# Best Cochlear Locations for Delivering Interaural Timing Cues in Electric Hearing

**DOI:** 10.1101/2024.12.27.627652

**Authors:** Agudemu Borjigin, Stephen R. Dennison, Tanvi Thakkar, Alan Kan, Ruth Y. Litovsky

## Abstract

Growing numbers of children and adults who are deaf are eligible to receive cochlear implants (CI), which provide access to everyday sound. CIs in both ears (bilateral CIs or BiCIs) are becoming standard of care in many countries. However, their effectiveness is limited because they do not adequately restore the acoustic cues essential for sound localization, particularly interaural time differences (ITDs) at low frequencies. The cochlea, the auditory sensory organ, typically transmits ITDs more effectively at the apical region, which is specifically "tuned" to low frequencies. We hypothesized that effective restoration of robust ITD perception through electrical stimulation with BiCIs depends on targeting cochlear locations that transmit information most effectively. Importantly, we show that these locations can occur anywhere along the cochlea, even on the opposite end of the frequency map from where ITD cues are most dominantly encoded in an acoustic hearing system.

## 1 Introduction

Human listeners rely on binaural hearing for everyday functions involving localization of sounds in the environment and segregation of sounds such as speech from background noise (Blauert, 1997; Middlebrooks & Green, 1991; Stecker & Gallun, 2012; Yost & Hafter, 1987). In typically hearing (TH) listeners, binaural hearing relies on the availability of acoustic cues, namely interaural level differences (ILDs) and interaural time differences (ITDs) (Rayleigh, 1909). ILDs and ITDs arise from the physical difference in the intensity and arrival time, respectively, of a sound between a listener’s ears. From a signal processing perspective, everyday sounds can be decomposed into amplitude and frequency modulations which are commonly referred to as the temporal envelope and temporal fine structure (TFS) in the field of hearing science. However, the distinction between the envelope and TFS becomes less clear for sounds with low-frequency content (Hilbert, 1906). TH listeners are more sensitive to ILDs in the high-frequency sounds (above 2 kHz) and to ITDs in the low-frequency sounds: whether in the TFS of low-frequency or in the slow envelope modulations of high-frequency TFS (Macpherson & Middlebrooks, 2002; Smith et al., 2002).

Cochlear implants (CIs) are electronic devices that provide access to sound to people with severe-to-profound deafness. In recent decades, bilateral CIs (BiCIs) have been clinically adopted to provide input to both ears, with one intention being the potential restoration of access to ILDs and ITDs (Brown & Balkany, 2007; Kan & Litovsky, 2015; van Hoesel & Tyler, 2003). Access to binaural cues can increase sound localization accuracy (Grantham et al., 2007; R. Litovsky et al., 2006; van Hoesel & Tyler, 2003), improve speech-in-noise understanding (R. Y. Litovsky et al., 2009; van Hoesel & Tyler, 2003) and reduce listening effort (K. C. Hughes & Galvin, 2013). However, listeners with BiCIs do not enjoy the same level of excellent sound localization (Anderson et al., 2022; Beijen et al., 2007; R. Litovsky et al., 2006; Verschuur et al., 2005; Zheng et al., 2015) or speech understanding in noise (R. Y. Litovsky, 2012; R. Y. Litovsky et al., 2009; Loizou et al., 2009; Ricketts et al., 2006) as TH listeners, with highly variable outcomes. Challenges with spatial hearing and speech-in-noise perception significantly reduce effectiveness of communications in professional and social interactions. For TH listeners, when both ILDs and ITDs are present, as in the case of a wideband sound common in everyday hearing, ITDs available in low-frequency TFS are generally prioritized over ILDs. Envelope ITDs (i.e., ITDs in the envelope of high-frequency TFS) contribute minimally, especially in the absence of noise interference (Blauert, 1997; Dennison et al., 2023; Jones et al., 2014; Macpherson & Middlebrooks, 2002; Middlebrooks & Green, 1991; Wightman & Kistler, 1992). For BiCIs, ILD cues are the primary binaural cue available (Aronoff et al., 2010; Grantham et al., 2007), in part because ITD cues are not preserved (Kan & Litovsky, 2015; van Hoesel & Tyler, 2003).

There are multiple reasons for weak or absent ITDs in BICIs. First, CI speech processors in the two ears are not synchronized. Thus, sounds reaching the microphones from locations in space can have an unwanted across-ear delay of hundreds of milliseconds (Dennison et al., 2022). This delay is highly problematic considering that the maximum ecologically relevant ITD is less than one millisecond (∼750 us) (Blauert, 1997; Kuhn, 1977; Middlebrooks & Green, 1991; Moller et al., 1995; R.S, 1907). Second, low-frequency TFS is not encoded in most clinically available CI sound coding strategies, which typically only extract the temporal envelope from each frequency channel (note that CIs typically decompose sounds into 12-22 frequency channels). Electrical pulse trains of high stimulation rate (∼1000 pulses per second; pps) are necessary to adequately sample and represent sound envelopes (Loizou et al., 2000). Indeed, CI listeners’ speech intelligibility has been shown to be better with high-vs. low-rate stimulation (Friesen et al., 2005; Loizou et al., 2000). However, just as TH listeners being more sensitive to ITDs in low-frequency sounds, BiCI listeners’ sensitivity to ITD has shown to be better at low stimulation rates (Anderson et al., 2019; Kan & Litovsky, 2015; Laback et al., 2007, 2015a; van Hoesel et al., 2009; van Hoesel & Tyler, 2003). The competing constraints of higher stimulation rates for speech understanding and lower stimulation rates for the delivery of ITD cues have yet to be reconciled. MED-EL is the only CI manufacturer that reports an attempt to include low-rate stimulation in their FSP and FS4 strategies (Riss et al., 2014). However, these commercial strategies have not shown a consistent benefit in spatial hearing outcomes (Ausili et al., 2020; Zirn et al., 2016), likely because MED-EL processors, like other devices, are not synchronized across the ears. The inconsistent improvement with unsynchronized processors suggests that lack of synchronization is preventing faithful delivery of ITD cues at low rates to listeners.

Our lab has taken a deliberate approach using synchronized research processors to investigate how low-frequency ITDs can be restored to BiCI users with the important goal of not sacrificing speech intelligibility. Our unique approach harnesses a novel speech coding strategy that aims to utilize “mixed rates” whereby selected pairs of electrodes in the two ears receive either low- or high-rate stimulation. The coding strategy, which is run on a bilaterally-synchronized processor, calculates an estimate of the ITD at the microphones and explicitly encodes a timing delay on the low-rate electrodes. This approach is a notable paradigm shift relative to today’s clinically fitted bilateral CI processors, which are not only unsynchronized, but stimulate all electrodes at fixed, high rates that obliterate the possibility of preserving low-frequency ITDs. Studies on our mixed rate strategy to date have demonstrated success in restoring BiCI listeners’ sensitivity to ITDs (Churchill et al., 2014; Thakkar et al., 2018, 2023), while also maintaining speech intelligibility (Churchill et al., 2014; Dennison et al., 2024).

While this finding occurs on a group level, large variability across patients indicates that mixed rate strategies only hold promise if we can further advance a more personalized medicine approach. This approach should take into consideration the impact of auditory deprivation on sensitivity to ITDs at different locations along the cochlea in the two ears. We base this premise on our work showing that individuals with earlier onset of deafness, i.e., auditory deprivation starting earlier in life, have poorer sensitivity to ITDs with low-rate stimulation (R. Y. Litovsky et al., 2010; Thakkar et al., 2020). While sensitivity to ITDs is vulnerable to auditory deprivation, there seems to be less impact on ILDs or ITDs in the envelopes of high-rate stimulation (R. Y. Litovsky et al., 2010; Thakkar et al., 2020; van Hoesel et al., 2009). Furthermore, amongst individuals whose deafness occurred later in life, there is variability in sensitivity to ITDs along the electrode array, suggesting that some places have sustained loss of neural elements or neural degeneration more than others (Anderson et al., 2023; R. Y. Litovsky et al., 2010; Thakkar et al., 2020). This issue is pertinent not only to human auditory neuroscience and the effects of auditory deprivation on binaural signal processing but also carries implications for clinical intervention.

The present study aims to understand how electrical stimulation with BiCIs can provide binaural benefits using a more personalized medicine approach. Prior studies tested all participants with the same set of mixed rate strategies; thus, all participants were stimulated with low-rate ITDs at the same pre-determined locations along the electrode arrays. A more personalized medicine approach potentially provides optimal encoding of ITD cues by taking advantage of the fact that each BiCI patient has a "best" location along the electrode array where ITD sensitivity is greatest. Hence, we hypothesized that benefits from a mixed-rate stimulation strategy will be maximized when these targeted “best” places are utilized for the delivery of low-rate ITD information, as compared to when the “worst” places are targeted.

The fact that the best performance with ITDs can potentially occur with low-rate stimulation presented anywhere along the cochlea, encourages a rethinking of how electric hearing can be utilized differently than acoustic hearing. The acoustic system relies on low-frequency information at the apical regions of the cochlea to promote best sensitivity to ITDs in the TFS. But in the electrical system, sensitivity to ITDs in low-rate stimulation has the potential to be achieved through stimulation anywhere along the cochlear electrode arrays. Some studies, such as (Best et al., 2011; Egger et al., 2014; Kan et al., 2013; Kan & Litovsky, 2015), have shown generally worse ITD sensitivity towards the apical-most place, while others such as van Hoesel et al., 2009 have shown an opposite trend, with worse ITD sensitivity measured with a basal electrode pair. Our current study was designed to investigate the extent to which variability in ITD sensitivity is observed across a group of BiCI users, in order to advance knowledge about which locations along the electrode array produce best sensitivity for each patient. Individualized information about the “best ITD place” was deemed necessary to test the hypothesis that, when multiple electrodes are activated in each ear in a mixed-rate strategy, we expected greater improvement from assigning low-rate stimulation to the electrode pair with the best ITD sensitivity than to the pair with the worst ITD sensitivity. Because multiple electrodes are a prerequisite for speech understanding (Fishman et al., 1997; Holmes et al., 1987), our multi-electrode stimulation study is a necessary step towards ultimately being able to preserve both ITD sensitivity for better sound localization and preserving speech information. This marks a pivotal move towards personalized medicine in the programming of BiCIs.

Each participant was tested with four strategies in total (please see *Figure 6* for illustration). Two of them were personalized mixed rate strategies, where the low-rate stimulation for ITDs were either assigned to a single electrode pair with the “best" or “worst" ITD sensitivity for the participant. To determine the location of the “best” and “worst” pair of electrodes, ITD sensitivity was first measured for each participant individually via a task of detecting the just noticeable difference (JND) in ITDs along the electrode array. The use of only one electrode pair (“best” or “worst”) for low-rate stimulation is intended to minimize the potential negative impact on speech understanding. We have previously shown that even allocating one electrode pair for low rates has the potential to improve ITD sensitivity in a mixed-rate strategy (Thakkar et al., 2018). Here we advanced a critical step towards determining the importance of allocating low rates not just to any place along the cochlea but to deterministically find the ideal location for each BiCI patient. Our approach was to compare stimulation strategies with the “best" or “worse” single low-rate channels to an "Interleaved" strategy, where every other channel gets low-rate stimulation, and a control condition, a clinical-like “All-high” strategy without any low-rate stimulation. All four strategies implemented for each participant contain the same set of 10 pairs of electrodes, which are roughly evenly spaced along the electrode array. To test our hypothesis, these four stimulation conditions were evaluated using a lateralization task, where participants reported perceived intracranial location of a stimulus, for a range of ITD values spanning the physiologically relevant range across the head. While the ITD JND task provides critical information about the variation in sensitivity to ITDs along the cochlea at a single-electrode level, the lateralization task with multi-electrode stimulation is more akin to real-world needs for localizing sounds in space. Previous studies on mixed rate strategies mainly used controlled, non-speech stimuli like complex tones (Thakkar et al., 2018, 2023), except for the recent work by Dennison et al., 2024. Here, we assessed the mixed rate strategies using a synthetic complex of tones and real speech stimuli. The speech stimuli provide greater temporal and spectral modulations and more realistic estimation of performance for everyday sounds. The comparison of synthetic and real stimuli allows us to understand the impact of real-world sounds on our mixed-rate strategy at an individual level.

## 2 Results

### 2.1 Measurement of ITD sensitivity along the cochlea

To measure ITD JNDs, each stimulation trial consisted of 2 stimulus intervals where each interval contained ITDs of the same magnitude but the leading ear is different (right-left or left-right). We aimed to measure the smallest ITD with which a participant could perceptually discriminate between a change in ITD. We measured ITD JNDs for fourteen BiCI participants. Figure 1 shows the ITD JNDs measured at five locations along the cochlea for each participant. It can be seen that the “best” and “worst” place for ITD JNDs can vary between individuals. Note that for some individuals, the difference in ITD JNDs between the best and second-best electrode pair locations is small (e.g., the ITD JNDs at basal and mid locations for IBF are nearly identical). The “best” and “worst” ITD JNDs for all participants are shown in Figure 1, with the location of the electrode pairs denoted by different symbols. Note that the “best” and “worst” values are statistically different for each individual. The differences are statistically significant at the p < 0.0001 level, based on a z-test and confidence interval estimates from bootstrap samples. A linear mixed effect model of the ITD JNDs with the location of the electrode pairs showed that there was no statistically significant predictive contribution from the place of stimulation (*F (*1,3) = 0.78, p = 0.513), confirming the fact that sensitivity to ITDs through low-rate stimulation can be achieved anywhere along the cochlear electrode array.

**Figure 1.**
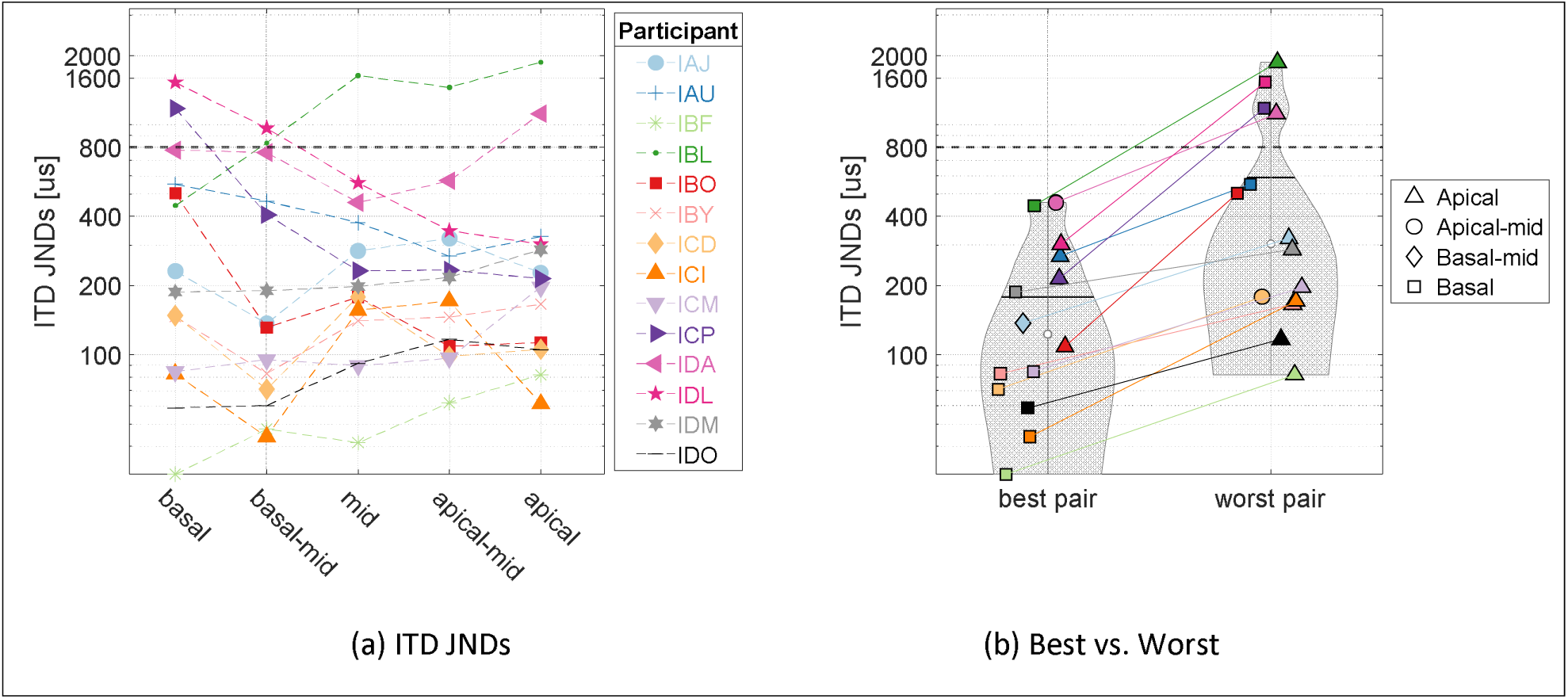
(a) ITD JNDs measured at 5 locations along the electrode array for all participants in this study (n = 14). The dashed line at 800 µs indicates the maximum ecologically relevant ITD for an adult human head. (b) Violin plot of the best vs. worst ITD JNDs for each individual (n = 14), with the location of electrode pairs labeled. This figure shares the same color code as panel a.

### 2.2 Comparing lateralization of sounds with mixed rates strategies

To test our hypothesis that a personalized strategy where low-rate stimulation is delivered to the places of best ITD sensitivity could maximize the benefit from a mixed rate strategy, the fourteen participants were tested on a perceptual lateralization task where they indicated the perceived intracranial location of a range of static ITDs (0, ± 200, ± 400, and ± 800 *µs*). The task was conducted with both synthetic (complex tone) and speech (CNC word) stimuli for four different strategies. The reported perceived locations were fitted with a psychometric function for each strategy, as shown in Figure 2. For subsequent analyses, we used the lateralization range from the fitted function as our metric, defined as the difference between the leftmost and rightmost locations. *Figure 3* shows the lateralization range data for complex tones (panel a) and CNC words (panel b). On a population level, the Interleaved mixed rate strategy resulted in largest ranges, out of all three mixed rate strategies (see Table 1 for stats). The Best mixed rate strategy led to larger lateralization than the All-high control. More importantly, the Best mixed rate strategy resulted in larger ranges, i.e., better performance than the Worst mixed rate strategies with the complex tone stimuli, as hypothesized. These observations are supported by the significant main effect of stimulation strategy on lateralization range (*F (*3,98) = 52.33, p *<* 0.001). Note that the comparison of the quantiles from model residuals and a sample normal distribution verified that the residuals follow a normal distribution. The residuals also passed the Shapiro-Wilk’s test for normality (*F (*13,98) = 1.37, p = 0.19). The residuals of the model passed Levene’s test for homogeneity of variance (grouped by strategy, *F (*3,108) = .74, p = 0.16; grouped by stimulus, *F (*1,110) = 1.10, p = 0.30).

**Figure 2.**
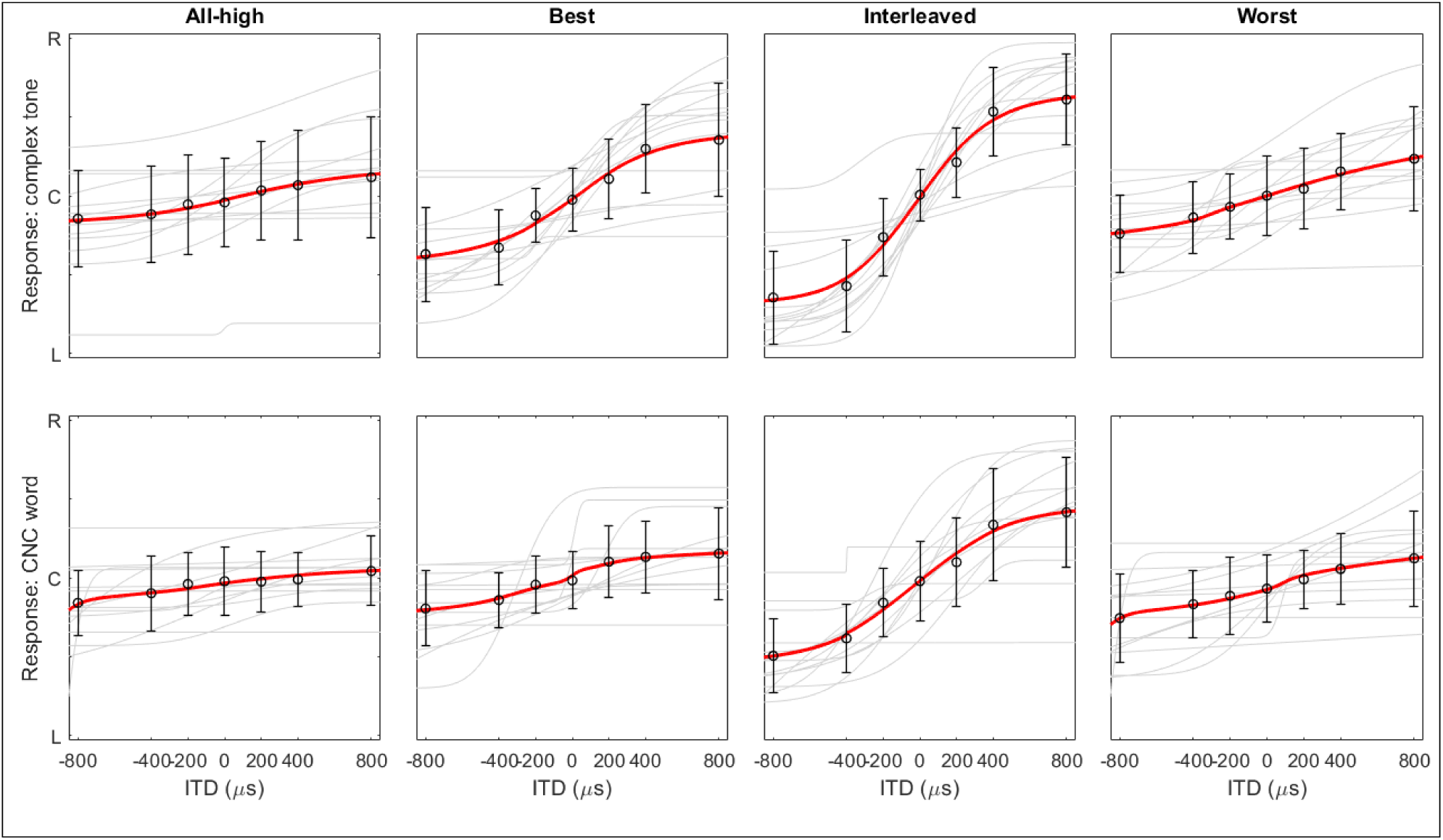
*Lateralization with complex tones (top row) and CNC words (bottom row). The light grey lines are the fitted psychometric functions for individual participants. The circles at each ITD are the group means of the fitted values while the error bar is 1 standard deviation. The red lines are the fit of group means. R, C, and L on the y axis represent right, center, and left and maps to ITDs of 800, 0, and –800 µs, respectively. For the fitted curves from each individual showing all data points, please see* Figure 7 *and* Figure 8.

**Figure 3.**
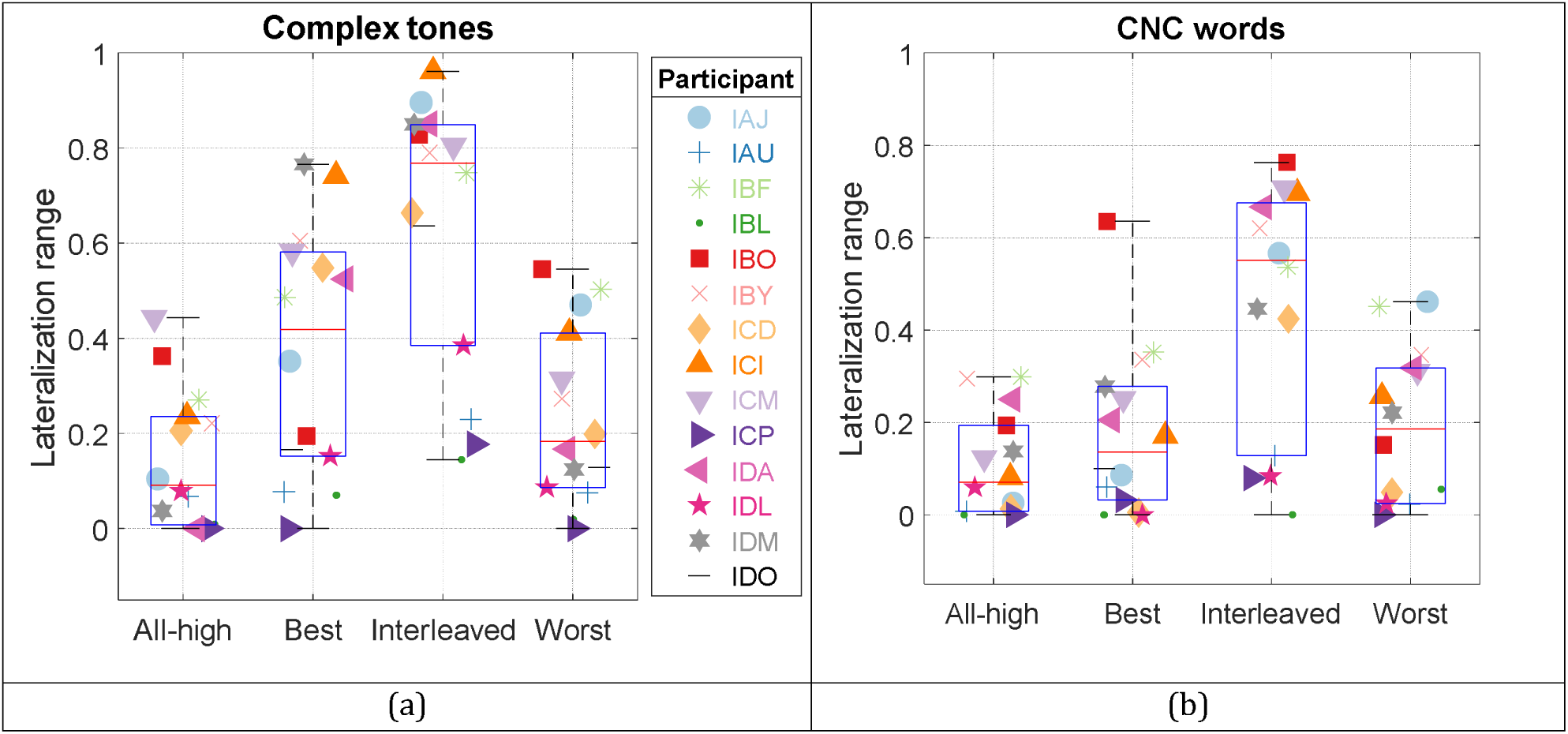
(a) Lateralization range for complex tone stimulus, plotted by processing condition, (b) same as (a) but with CNC word stimulus. The lateralization range of 0 and 1 corresponds to not perceiving a change in intercranial location with ITDs and being able to use ITDs for the full range of lateralized perception, respectively.

**Table 1.**
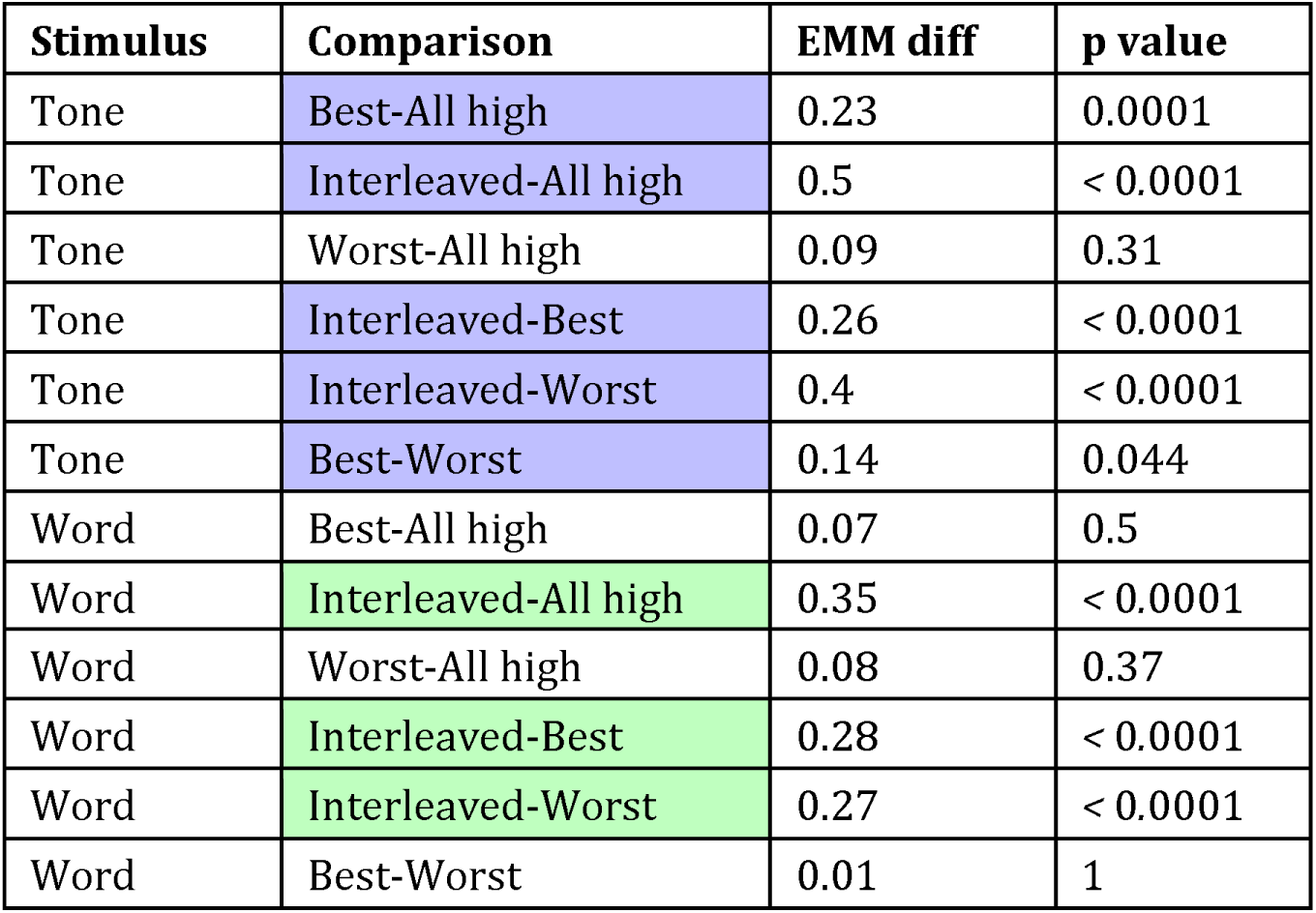
Post hoc comparisons were evaluated with Estimated Marginal Means (EMM) for processing strategies with different stimulus types. Statistically significant comparisons are shaded in color (purple for complex tones, green for CNC words). We used the Bonferroni correction for a family with 6 comparisons.

Additionally, lateralization range with the complex tones was significantly larger than with CNC words (EMM diff (tone - word) = 0.12, p < 0.001; EMM: Estimated Marginal Means), reflected by the significant main effect of stimulus type (*F (*1,98) = 21.23, p *<* 0.001) on lateralization range. With CNC word stimuli, the benefit of Best vs. Worst mixed rate strategy does not hold (EMM diff (best - worst) = -0.01, p = 1). The benefit from the Interleaved mixed rate strategy was also reduced with CNC word stimuli (EMM diff (tone – word) = 0.18, p = 0.0007), despite the statistical significance for all comparisons with other strategies (Table 1). Most participants did not benefit from their Best mixed rate strategy with speech stimuli. This was probably due to speech stimuli being more spectra-temporally sparse at the basal, high-frequency channels. Despite our best efforts to ensure stimulation across all channels by selecting CNC words that are more broadband, higher frequency channels still provided less stimulation than lower frequency channels (see *Figure 4*). Note that for most participants (8 out of 14), the best ITD JNDs were measured at basal, high-frequency electrode pair (see Figure 1). The lack of high frequency energy in the CNC words would mean that the low-rate stimulation at basal channels in the Best mixed rate strategy might not have been long enough to provide sufficient encoding of ITDs for lateralization in most participants, which could explain the lack of benefit. In contrast, participant IBO, whose best ITD JND was measured at the apical electrode pair, showed substantial benefit with the Best mixed rate strategy (see *Figure 3* for the lateralization benefit with CNC words and *Figure 4* for an example electrodogram for CNC word stimuli for this participant).

**Figure 4.**
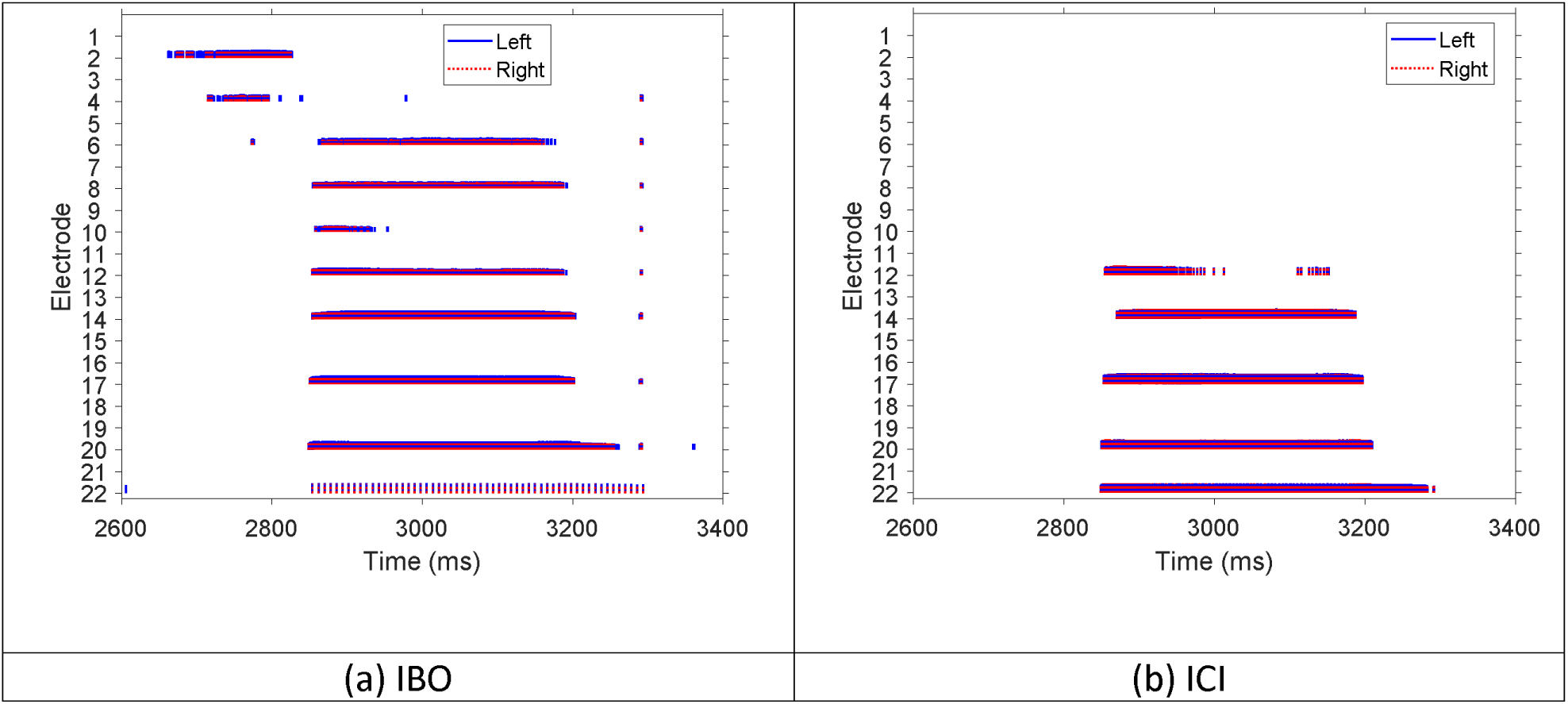
Electrodograms of the Best mixed rate strategy. Electrode 1 and electrode 22 correspond to the highest and lowest frequency channel, respectively. Pulses on the right and left side are shown in red and blue, respectively. Stimuli: CNC word. IBO’s best ITD sensitivity was obtained with the apical-most channel (i.e., low frequency) hence low-rate stimulation was sent to electrode 22 (i.e., see the sparse stimulation pattern at electrode 22). ICI’s best-ITD-sensitivity or low-rate channel is at the basal location. But there was no stimulation energy in high-frequency channels in this example.

### 2.3 The effects of place for low-rate stimulation

Figure 5 illustrates lateralization range with the Best mixed rate strategy when using the complex tone stimuli, categorized by the stimulation site that received the low-rate stimulation (i.e., the electrode pair with the best ITD JND). Overall, lateralization range is better with low-rate stimulation being sent to the basal electrode pair than apical electrode pair (marginal statistical significance based on Mann-Whitney test: *z* = −1.97, p = 0.048; non-parametric test was used since data do not follow normal distribution). Poor lateralization with apical electrode pair receiving low-rate stimulation can be explained by poor ITD sensitivity: three of the four participants (IAU, ICP, and IDL)’s best ITD JNDs are among the highest (327, 214, 303 µs, respectively). However, the remaining one of the four participants, IBO, showed good sensitivity to ITD with their best pair of electrodes (i.e., apical; JND = 113 µs), but still had poor lateralization when the low-rate stimulation was sent to their apical pair of electrodes. In contrast, IBO’s lateralization was much better when the low-rate stimulation was sent to the basal electrode pair, although it led to the worst sensitivity to ITD for IBO (JND = 504 µs, see Figure 3 for IBO’s lateralization with the Worst mixed rate strategy). As mentioned earlier, previous studies from our lab examined mixed rate strategies with various channel allocations for low-rate stimulation, but these were assessed at a population level without using ITD sensitivity measures for guiding channel selection as in the current study. Nevertheless, we extracted lateralization range data (Thakkar et al., 2023) and ITD JND discrimination thresholds (Thakkar et al., 2018) with similar strategies as in current study, plotted with permission in Figure 5 and Figure 5, respectively. These data show a similar pattern: mixed rate strategy with low-rate stimulation allocated at apical electrode locations led to poorer performance. These following participants from the current study participated in both prior studies in Figure 5 b and c: IBF, IBY, ICD, ICI, ICP. In both prior studies, all these five participants included in the current study showed greater benefits from low-rate stimulation with a basal electrode pair compared to an apical electrode pair. Note that although the strategies are similar, these mixed rate strategies in earlier studies contained only five channels, which was extended to ten in this study. In addition, these earlier studies used direct stimulation, with exact control over every aspect of the stimulus. The current study, on the other hand, manipulated the input signal using different processing strategies, which is more similar to what happens with clinical processors in real-world listening.

**Figure 5.**
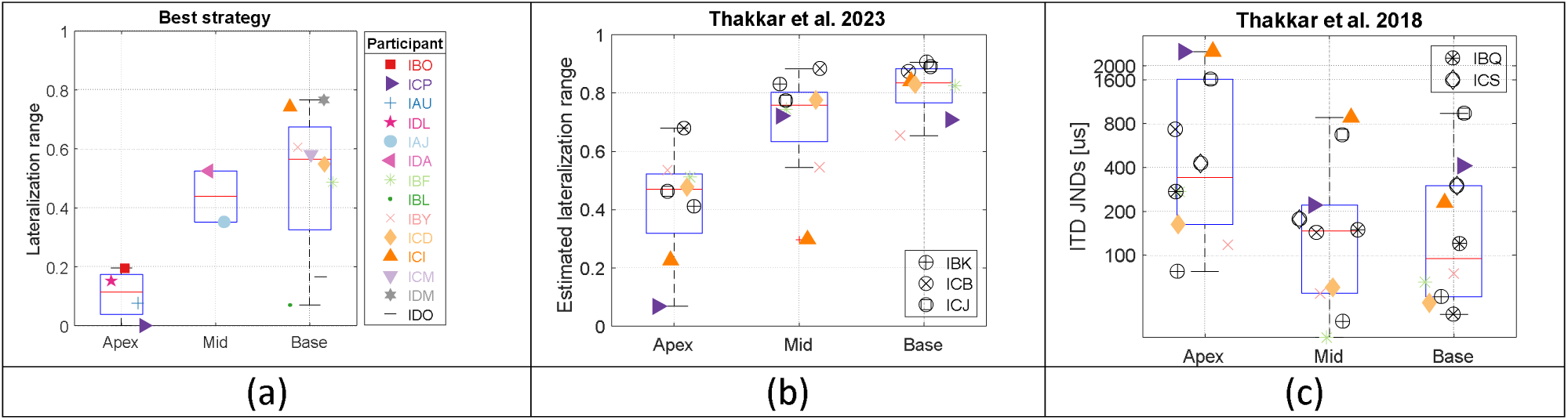
(a) Lateralization range measured with the Best mixed rate strategy (stimuli: complex tone), grouped by the location of the low-rate stimulation in the mixed rate strategy. (b) Lateralization data from similar strategies published previously from the lab (Thakkar **et al.**, 2023). Note that only the participants who did not participate in the current study were labeled. All three types of mixed rate strategies (i.e., with low-rate stimulation delivered to the single electrode pair at the apical, mid, and basal regions, respectively) were tested on each participant in this prior study (i.e., without customizing channel selection for low-rate stimulation). c) ITD JNDs measured with the same set of 3 mixed rate strategies on all participants from Thakkar **et al.**, 2018. Note that the participants who did not participate in the current study or the study in panel b were labeled. Data from panels b and c were re-plotted here with permission from Thakkar **et al.**, 2023, copyright 2023, Acoustical Society of America (doi) and Thakkar **et al.**, 2018; copyright 2018, Acoustical Society of America (doi), respectively.

## 3 Discussion

This study aims to understand how electrical stimulation with BiCIs can deliver binaural benefits through a more personalized medicine approach. We hypothesized that ITD sensitivity varies along the electrode array per each individual and that targeting regions with the best ITD sensitivity could optimize performance when providing low-rate ITD cues. Based on this hypothesis, we predicted that a mixed rate strategy targeting the electrode with the best ITD sensitivity would outperform one targeting the worst ITD sensitivity. Likewise, we expected that the mixed rate strategy with a single “worst” channel would result in a similar performance to a high-rate only coding strategy without any low-rate ITD cues. These are indeed what we found with the complex tone stimuli.

Previous mixed rate studies typically only used controlled, non-speech stimuli, such as single- or multiple-electrode pulse trains (Thakkar et al., 2018, 2023), except for a few studies such as Churchill et al., 2014 and the most recent work from Dennison et al., 2024. We therefore also evaluated the mixed rate strategies with speech stimuli, which offered more temporal and spectral modulations. The benefits to lateralization from the Best mixed rate strategy with a simple stimulus did not hold with speech stimuli. For both stimulus types, the Interleaved mixed rate strategy outperformed all other conditions. This suggests that more than one low-rate channel might be necessary for more redundancy, especially when the stimuli are more spectral-temporally dynamic such as speech (Ding et al., 2017; Elliott & Theunissen, 2009). The ITD cue was perhaps not consistently present due to the temporal modulations inherent to speech or lack of stimulation energy at high-frequency channels (in case high-frequency, basal channels were selected for low-rate stimulation in the “Best” strategy). Considering the best ITD sensitivity has been usually observed with basal-most channels, this factor should be considered when selecting channels for low-rate stimulation based on ITD sensitivity. Nevertheless, real-world communication typically involves listening to running speech instead of single-syllable words such as those used here. A running speech that consists of several words might offer more opportunities for stimulation at basal, high-frequency channels. Another thing to note, however, is that amplitude modulations in speech stimuli introduce envelope-based ITD cues in addition to the pulse-timing or TFS ITD cues. Dennison et al., 2024 observed the tendency of lateralization performance being better with pulse-timing ITDs only compared to when both envelope and pulse-timing ITDs were present (not statistically significant). This suggests a potential conflict when listeners are exposed to stimuli containing both types of ITD cues. The mixed rate strategies tested in their study were like the Interleaved strategy in the current study. Too many low-rate channels might have contributed to this conflict. As mentioned, too many low-rate channels can be problematic also in terms of compromising speech intelligibility (Friesen et al., 2005; Loizou et al., 2000). In summary, an optimal number of channels reserved for low-rate stimulation, between single and half of all channels, might be key to maximizing the benefits of a mixed rate strategy, especially with speech signals.

Interestingly, we also observed that more participants had their best ITD sensitivity measured with a single basal pair than with a single apical pair (Figure 1). This is consistent with several previous studies (Best et al., 2011; Egger et al., 2014; Kan et al., 2013; Thakkar et al., 2020), despite differences in task parameters. We used stimulation rate of 125 pps instead of 100 pps as in all four cited studies; we also directly paired electrodes by their numbers (e.g., electrode 4-4), without an additional pitch-matching procedure. This finding, i.e., more participants having their best ITD sensitivity measured with basal electrode pair, indicates that electrical stimulation presents a different pattern than acoustic stimulation in a TH auditory system, which is that TFS-based ITD processing is confined to the low-frequency, apical regions of the basilar membrane (Brughera et al., 2013; J. W. Hughes, 1940). The auditory system likely cannot phase lock to the fine structure of sounds above 1600 Hz and diminishes as frequency increases (Brughera et al., 2013; J. W. Hughes, 1940; Johnson, 1980; Palmer & Russell, 1986; Verschooten et al., 2019). MED-EL, the only CI manufacturer that claims to implement a strategy aimed at encoding TFS, assigns low-rate stimulation to low-frequency, apical channels [FS-strategy family, launched in 2006 (Riss et al., 2014)]. However, the spatial and speech-in-noise hearing benefits of this approach have been mixed: while studies have shown benefits in spatial hearing (Fischer et al., 2021) and overall speech-in-noise hearing (Lorens et al., 2010; Vermeire et al., 2010), others, such as Zirn et al., 2016 do not. These mixed results could be explained by the relatively poorer sensitivity to ITDs with the low-frequency, apical stimulation channels, as shown in this study. Participant IBO is one of the exceptions, whose best ITD sensitivity was measured with apical electrode pair. However, lateralization performance with complex tone was worse with low-rate stimulation being delivered at the apex than base (worst ITD sensitivity for IBO) in a mixed rate strategy. This suggests that the benefits of mixed rate strategy in lateralization tend to be greater when the low-rate stimulation is assigned at the basal electrode pairs, regardless of their ITD sensitivity. This pattern was also shown by similar conditions tested in a couple of previous studies from the lab (Thakkar et al., 2018, 2023) (see Figure 5). To explain why apical electrodes tend to have worse ITD sensitivity, we argue that neural stimulation is less selective at the apical region due to “cross-turn” stimulation (Briaire & Frijns, 2006), and that electrical stimulation properties are not well-tuned to neural structures that typically encode ITD information (Laback et al., 2015b). The assignment of low-rate stimulation to basal electrode pairs has the benefit of minimizing the negative impact from introducing low-rate stimulation on speech understanding (Friesen et al., 2005; Loizou et al., 2000), considering most of the speech energy is in the range from 200 to 3500 Hz, not so much in the basal-most channels (Sobolewski, 2003).

In summary, as hypothesized, while it is important to target the electrode location with the best ITD sensitivity for low-rate stimulation, the electrode location with the best ITD sensitivity seems to be similar across participants (basal electrode pairs for most participants). For maximum benefit with speech stimuli, more than one channel should be reserved for low-rate stimulation. If two channels were to be selected for low-rate stimulation, one would still be at the basal locations, where ITD sensitivity is likely to be the best, although speech energy tends to be less in those channels. The other electrode location should avoid apical region, where ITD sensitivity may be poor. However, the mixed rate strategy might benefit from selecting a site where speech energy is typically more dominant (i.e., somewhat away from the basal most channel). These findings could inform the development of more personalized CI programming strategies, potentially leading to improved outcomes for BiCI users in both speech understanding and spatial hearing, thereby enhancing their overall quality of life. One limitation of this study is the controlled laboratory setting in which the ILD cues were minimized from the stimuli. Real-world listening conditions, where ILD and ITD cues coexist, might yield different outcomes. Like the potential conflicts between the envelope and pulse-timing ITD cues, an interaction might arise between ITD and ILD cues. Future research could explore the long-term adaptability of patients to mixed rate strategies in diverse auditory environments, potentially examining the interaction between ITD and ILD cues under dynamic listening conditions.

## 4 Materials and Methods

In this study, we constructed “best” and “worst” mixed rate strategies, where a single pair of electrodes with the best or worst ITD sensitivity was selected for low-rate stimulation, respectively. We conducted ITD discrimination tasks along the electrode array to evaluate ITD sensitivity at different locations. “Interleaved” mixed rate strategy, where every other channel received low-rate stimulation, was also created to investigate whether performance with a single-channel mixed rate strategy (Best mixed rate strategy) would be similar to performance with multi-channel mixed rate strategy. Three mixed rate strategies, along with a control strategy without any low-rate stimulation, were evaluated with a lateralization task.

### 4.1 Participants

Fourteen BiCI users participated in this study. Participants traveled to the University of Wisconsin-Madison for three days of testing. They were paid a stipend for their participation and all travel-related expenses were compensated. Participant demographics are displayed in *Table 2*. All participants had Cochlear Ltd. Implants (Sydney, Australia). All experimental procedures followed the regulations set by the National Institutes of Health and best practices for direct stimulation studies (R. Y. Litovsky et al., 2017), and were approved by the University of Wisconsin-Madison Health Science Institutional Review Board.

**Table 2.** Demographic and implant information for BiCI listeners. EVA: enlarged acoustic aqueduct; ISL: idiopathic sudden loss; SI: skull injury; H: hereditary; U: unknown; O: otosclerosis.

| ID | Sex | Age at testing (years) | Age at hearing loss (L, R) | Age at implantation (L, R) | Experience with BiCIs (years) | Bilateral hearing loss before BiCIs (years) | Etiology (L, R) |
| --- | --- | --- | --- | --- | --- | --- | --- |
| IAJ | Female | 78 | 5, 5 | 51, 58 | 20 | 53 | H, H |
| IAU | Male | 74 | 3, 3 | 50, 56 | 18 | 53 | H, H |
| IBF | Female | 72 | 38, 38 | 56, 54 | 16 | 18 | H, H |
| IBL | Female | 77 | 12, 12 | 54, 59 | 18 | 47 | U, U |
| IBO | Female | 58 | 23, 23 | 45, 42 | 13 | 22 | O, O |
| IBY | Female | 60 | 41, 41 | 43, 48 | 12 | 7 | U, U |
| ICD | Female | 65 | 3, 3 | 50, 44 | 15 | 47 | EVA, EVA |
| ICI | Female | 65 | 31, 31 | 50, 51 | 14 | 20 | U, U |
| ICM | Female | 70 | 23, 23 | 57, 58 | 12 | 35 | U, U |
| ICP | Male | 61 | 4, 4 | 46, 49 | 12 | 45 | U, U |
| IDA | Female | 57 | 8, 8 | 47, 46 | 10 | 39 | U, U |
| IDL | Female | 69 | 33, 33 | 62, 61 | 7 | 29 | U, U |
| IDM | Female | 46 | 5, 5 | 33, 35 | 11 | 30 | U, U |
| IDO | Male | 52 | 46, 38 | 46, 43 | 6 | 0 | SI, ISL |

### 4.2 Experimental Design and Statistical Analyses

#### 4.2.1 Experiment conditions

Four stimulation strategies were compared in this study (Figure 6, panel C visually summarizes each strategy): All-high, Interleaved mixed-rate, Best mixed-rate, and Worst mixed-rate. Each participant had the same set of ten electrodes activated for both ears for all four strategies. The All-high strategy used high-rate stimulation of 1000 pulses per second (pps) at all ten electrode pairs. In the Interleaved mixed rate strategy, every other electrode pair received low-rate stimulation of 125 pps. Both the Best and Worst mixed rate strategies had a single pair of electrodes stimulated at low rate (125 pps), while the remaining nine pairs of electrodes received high-rate stimulation of 1000 pps. For the Best mixed rate strategy, the low-rate stimulation was sent to the electrode pair with the lowest (i.e., best) ITD JND (see Figure 6, panel B: basal electrode pair 4-4). Accordingly, in the Worst mixed rate strategy, the low-rate stimulation was sent to the electrode pair with the highest (i.e., worst) ITD JND (see Figure 6, panel B: apical electrode pair 22-22). ITD JNDs were determined at each of those 5 low-rate electrode pairs with an ITD discrimination task (for details, see section 4.2.2). See Figure 6 panel B for an example set of ITD JND measurements at these 5 low-rate locations. Each processing strategy was implemented using custom MATLAB software written for the CCi-MOBILE, a bilaterally synchronized and portable CI research platform (Ghosh et al., 2022; Ghosh & Hansen, 2023). See (Dennison et al., 2024) for more details on how these stimulation strategies were implemented on the CCi-MOBILE.

**Figure 6.**
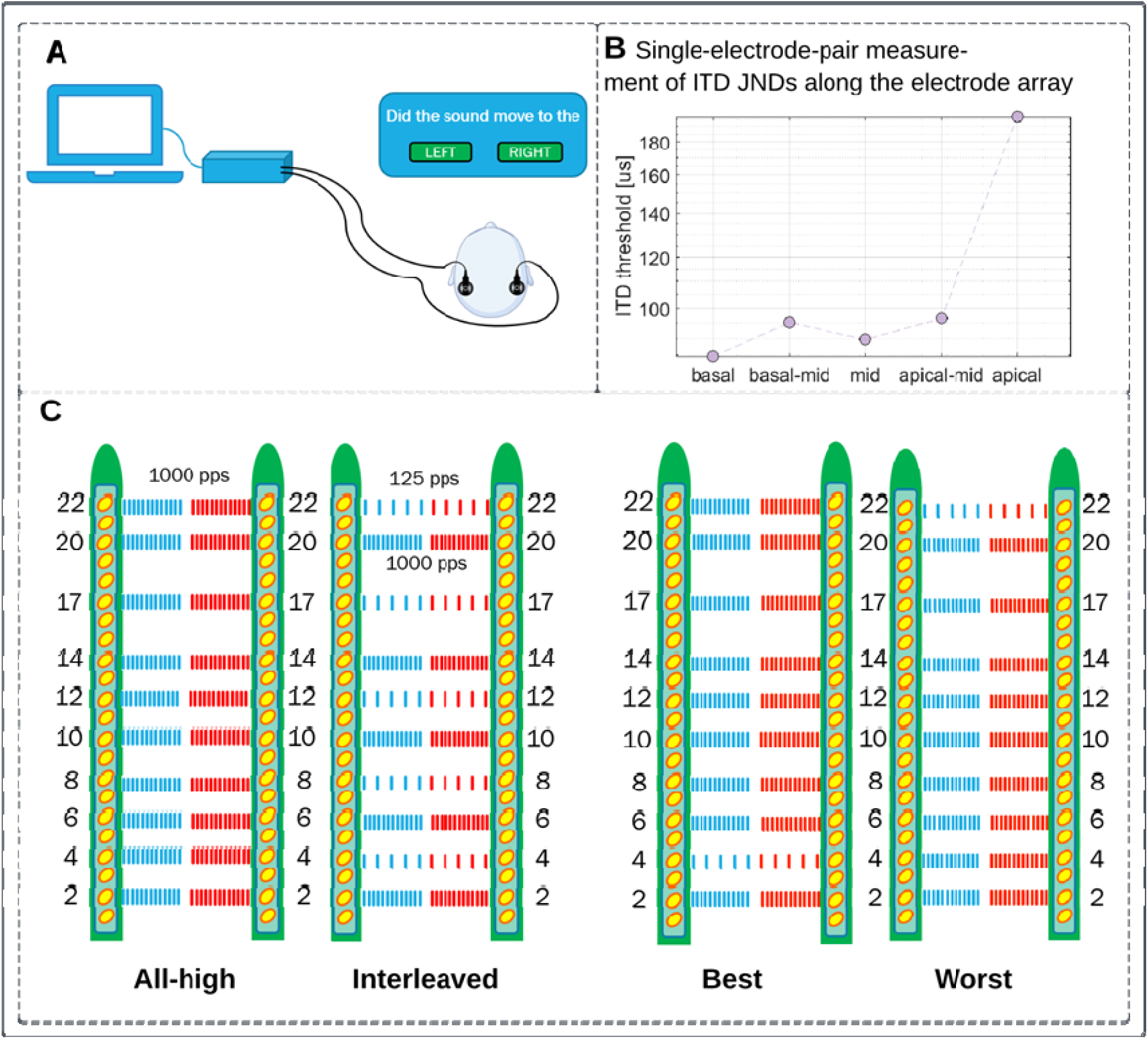
Stimulation strategies/conditions. A. Direct stimulation setup for ITD JND measurement to produce data shown in panel B. B. Five ITD JNDs measured with the low-rate electrode pairs as in the Interleaved condition in panel C: 4-4 (basal), 8-8 (basal mid), 12-12 (mid), 17-17 (apical-mid), 22-22 (apical). C. Four stimulation strategies/conditions evaluated in this study.

**Figure 7.**
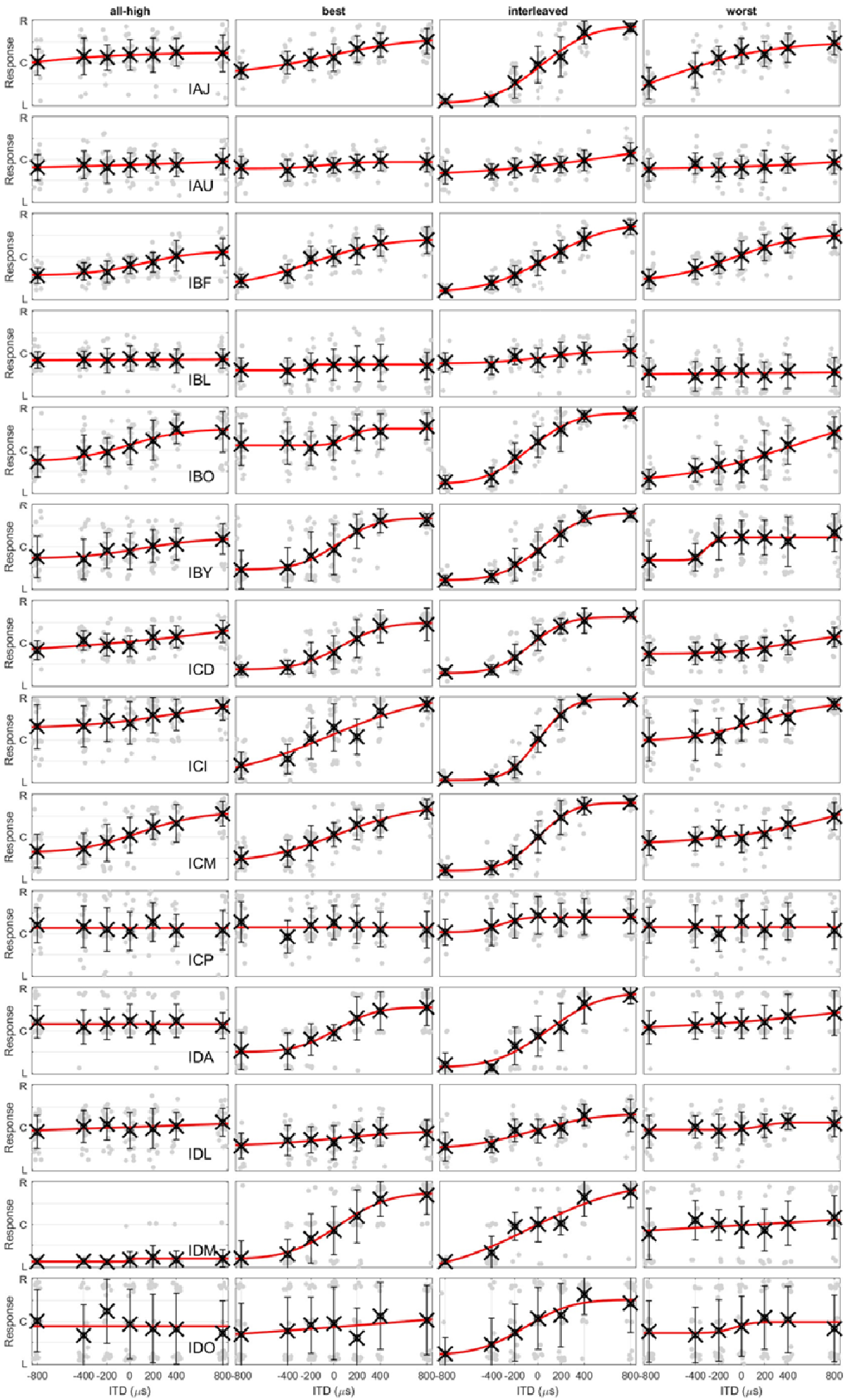
Lateralization with complex tones. Each row contains data with 4 strategies from an individual.

**Figure 8.**
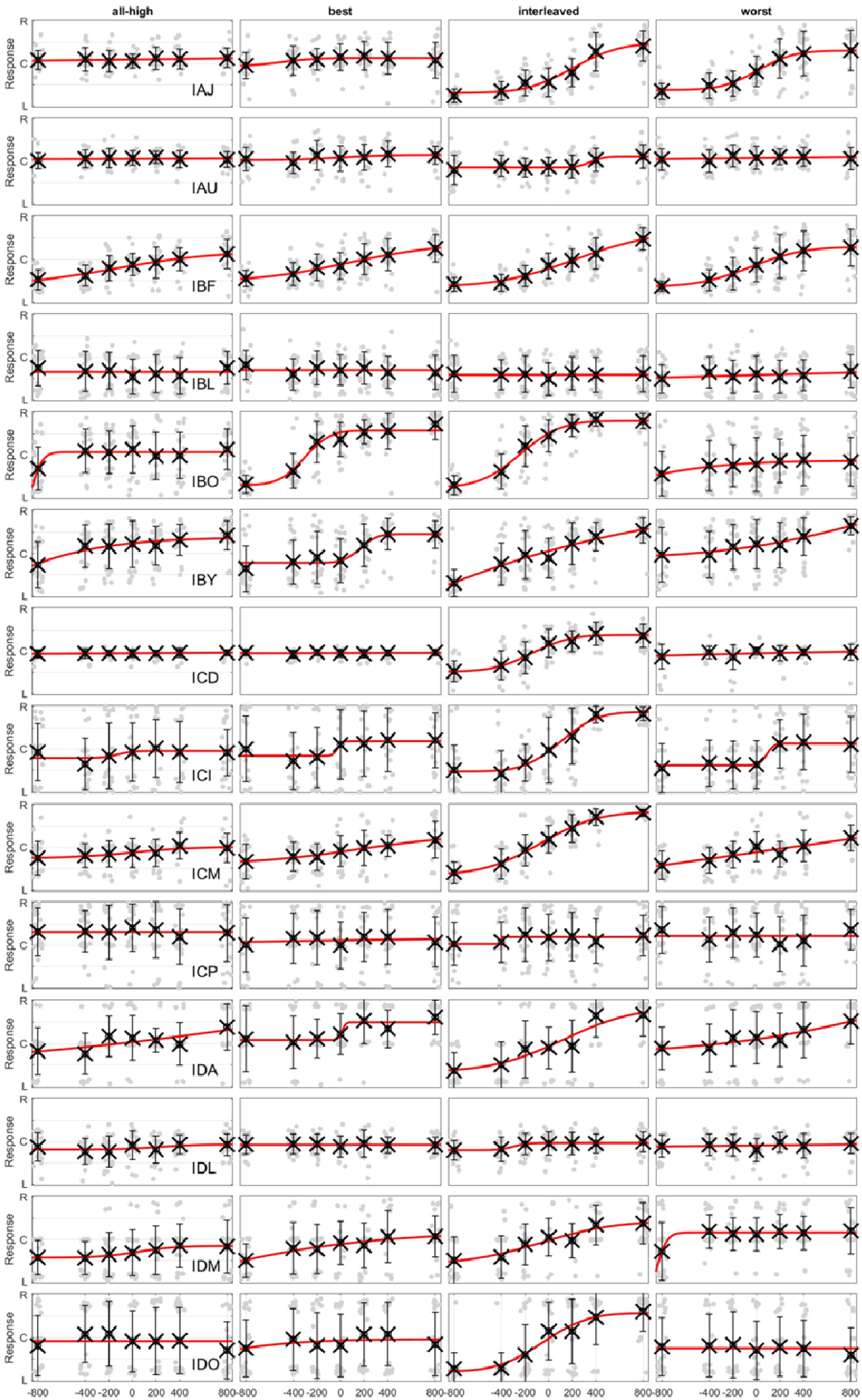
Lateralization with CNC words. Each row contains data with 4 strategies from an individual.

#### 4.2.2 Stimuli, procedure, and equipment

##### Devices

Loudness mapping, ITD discrimination (procedures described below) used the Nucleus Implant Communicator (NIC) libraries in MATLAB (Mathworks, Natick, MA) to communicate with the RF GeneratorXS (Cochlear, Sydney, NSW, Australia). Custom-written MATLAB (R2022b) software was used to create the testing interface, which generated and sent the stimuli directly to the participant’s implants. Although RF GeneratorXS can be used for lateralization with multi-electrode stimulation, we used the CCi-MOBILE because it allows for testing research strategies in real-time. This research is also partly funded by a grant on the CCi-MOBILE (NIDCD: R01-DC016839). Compared to relatively bulky RF GeneratorXS, the CCi-MOBILE is a much smaller and hence portable research platform, which is bilaterally synchronized, meaning that a single clock is used to drive two internal devices simultaneously (see Dennison et al., (2022) for a discussion on synchronized processors). We can use the CCi-MOBILE for simultaneous processing and stimulation of a pair of Cochlear internal implants via a computer running Microsoft Windows (V10) (Redmond, WA). The CCi-MOBILE has been demonstrated as a suitable platform for streaming binaural audio and studying the lateralization abilities of BiCI users (Dennison et al., 2023).

##### Loudness mapping

Prior to testing ITD discrimination and lateralization, threshold (T) and most comfortable (C) loudness levels were measured using custom MATLAB software with RF GeneratorXS. Mapping stimuli were 300 ms constant amplitude pulse trains at a rate of 125 or 1000 pps (depending on high or low-rate channels). Pulse widths matched each participants’ clinical setting. Inter-phase gap duration was set at 8 us. T and C levels were only remeasured for the ten electrodes selected for the stimulation strategies. Four maps (one for each stimulation strategy) were created for this study. Each map uses the same set of ten electrodes with default selection of electrodes 2, 4, 6, 8, 10, 12, 14, 17, 20, 22 for both sides. The selection of the electrodes was adjusted if there were any deactivated electrode in a participant’s clinical map. For example, if electrode 4 is deactivated in participant’s clinical map, we either choose electrode 3 or 5 instead. For the Interleaved mixed rate strategy, electrodes 4, 8, 12, 17, 22 were assigned as low-rate channels by default (see Figure 6, panel C, Interleaved condition). Following measurement, all electrodes were loudness balanced within each ear. To do so, multiple electrodes were stimulated at their C levels in sequence, first in groups of three adjacent electrodes, with overlapping electrode between two adjacent electrode groups, then groups of five adjacent electrodes. Loudness was also balanced across the ears by stimulating single electrode pairs simultaneously, making sure that the stimulation resulted in a centered intracranial percept. Note that the loudness was also balanced for each of the four stimulation strategies by adjusting the overall stimulation level for two sides when all electrodes were stimulated. Finally, overall loudness was balanced across 4 stimulation strategies.

##### ITD discrimination

ITD discrimination was tested with a 2-interval, 2-alternative forced-choice (2AFC) task. The stimulus in each interval was a 300-ms, constant amplitude pulse train at 125 pps, presented with a delay between the two ears. Inter-stimulus interval was set at 300 ms. The magnitude of the ITD was the same in both intervals, but the polarity was opposite between intervals. Listeners were asked to indicate the perceived direction of the second interval relative to the first. We used the method of constant stimuli to measure discrimination thresholds, with a default selection of ITDs: 50, 100, 200, 400, 800 us. If necessary, additional ITDs below 50 µs and/or above 800 µs were added to complete a psychometric function based on percent correct scores. To determine whether extra ITDs were needed, data collection was broken into many runs and the data was plotted after each run. The JND was estimated as the 75% point along the psychometric curve (Wichmann & Hill, 2001). The data were fitted using the psignift MATLAB package (version 2.5.6) (Kuss et al., 2005). Each ITD was presented 40 times to each electrode pair, with half right-leading (i.e., 20 times) and half left-leading. The order of presentation for ITDs of different magnitude and polarity was randomized. The ITD JND was measured at one electrode pair at a time. Note that the stimulation levels on two sides were adjusted to elicit a centered auditory image (i.e., C levels were balanced across ears, see procedure above), or in other words, ILD information was set to 0 dB. An initial training with feedback was provided before the formal data collection. Feedback was turned off during the task.

##### Lateralization

Lateralization stimuli were presented through the four research strategies (Figure 6) implemented on the CCi-MOBILE. Two different stimulus types were presented to listeners for lateralization: tone complexes and CNC words. Tone complexes were generated at a sampling rate of 96 kHz as acoustic wave files. Tone complexes were created by summing ten sinusoids with frequencies corresponding to the center frequencies of the ten bandpass filterbank channels (see Table 3 for details). Tone complexes were 300 ms in duration. Wave files for CNC words were previously recorded in our lab. Only female recordings were used to ensure that all channels, especially the high-frequency channels, were stimulated. This is important because some participants might have their best ITD sensitivity measured with basal electrode pairs, which leads to the selection of those channels for low-rate stimulation in their Best mixed rate strategy. Using different word for every trial introduced too much spectral variation from trial to trial, which might present challenge for interpreting the results. However, using only one CNC word across all trials led to boredom or distraction among participants during pilot testing. Therefore, only 5 different words (sob, can, sail, lash, voice) were used across all trials. All CNC recordings began with a “Ready” cue before the monosyllabic word was presented (this word “Ready”was also lateralized along with the CNC word).

**Table 3.** Frequency allocation table (FAT). Some participants used adjusted Electrode numbers but had identical frequency allocations. Cochlear device contains 22 intra-cochlear and 2 ground electrodes. The numbering convention for electrodes of these Cochlear devices is that 22 is the apical-most electrode and 1 is the basal-most electrode.

| Channel | Electrode | Cutoff frequency (Hz) |  |  |
| --- | --- | --- | --- | --- |
|  |  | Lower | Center | Upper |
| 1 | 2 | 6063 | 6501 | 6938 |
| 2 | 4 | 4688 | 5001 | 5313 |
| 3 | 6 | 3563 | 3813 | 4063 |
| 4 | 8 | 2688 | 2876 | 3063 |
| 5 | 10 | 2063 | 2188 | 2313 |
| 6 | 12 | 1563 | 1688 | 1813 |
| 7 | 14 | 1188 | 1251 | 1313 |
| 8 | 17 | 813 | 876 | 938 |
| 9 | 20 | 438 | 501 | 563 |
| 10 | 22 | 188 | 251 | 313 |

The lateralization task was completed on a Windows Surface computer (Microsoft, Redmond WA, USA; Intel(R) Core(TM) i7-1065G7 CPU @ 1.30GHz 1.50GHz, 16 GB RAM), using the method of constant stimuli in a single-interval paradigm. Listeners used a Graphical User Interface (GUI) to initiate presentation of each stimulus with a “Play” button. The GUI had a cartoon image of a face with a bar overlaid on top. The listener put a visual marker in the bar at the location where they perceived the sound to originate inside their own head. Each location on the bar in the GUI was converted into a value between –0.5 and 0.5, with negative and positive numbers indicating left and right locations, respectively. There was a button for repeating the stimulus, but listeners could repeat each trial only once. For each stimulus type, listeners were presented with ITDs of magnitude +/-200, +/-400, +/-800, and 0 µs. Five repetitions of each ITD were included in a single block, and the presentation order was randomized within a block. Within each stimulus type, the four strategies were tested with a 4x4 Latin Square block design to counterbalance order effects within the group (i.e., a total of 16 blocks or 4 blocks for each strategy for each stimulus type). The presentation order of the stimulus type was counterbalanced across listeners, with half completing lateralization trials with tone complexes first and the other half with CNC words first.

#### 4.2.3 Statistical Analyses

Statistical analysis was performed with RStudio (R version 4.3.1). To test the prediction that individuals vary in their ITD sensitivity at different locations along the electrode array, the ITD JNDs were fitted using linear mixed effect model (lme4 package, version 1.1.31) with electrode pair location being independent factor, and with random effects to account for the variability associated with participants: *model* = *lmer*(*ITD JNDs* ∼ *electrode locations* + 1|*participant*). To test the prediction that BiCI listeners would show better lateralization performance with the Best than with the Worst mixed rate strategy, we used a linear mixed effects model with lateralization range being the dependent factor and stimulation strategy being independent factor. We also included stimulus type as an independent factor to investigate the influence of stimulus type on the performance. For analyzing lateralization data, the raw data were first fitted with Nonlinear Least Squares (curve fitting function: fit) in MATLAB. We then extracted the difference between the top and bottom asymptotes as lateralization range. We also attempted analyzing the of slope on the lateralization curve. However, the lateralization curves are too variable across participants to standardize a slope analysis. To summarize, the lateralization range data were fitted using linear mixed effect model with stimulation strategy and stimulus type being independent factors, and with random effects to account for the variability associated with participants: *model* = *lmer*(*lateralization range* ∼ *stimulation strategy* ∗ *stimulus type* + 1|*participant*). The anova function (package car, version 3.1-2) was used to calculate the type-III sequential sum of squares for analyzing the predictive contribution from the independent factors and their interactions in the linear mixed effects model. The normality of the variance was inspected both visually by comparing the quantiles from model residuals and a sample normal distribution and conducting Shapiro-Wilk tests on the residuals. The homogeneity of the variance was inspected by conducting Levene’s test on the model residuals. We conducted post-hoc comparison analysis by using the emmeans (version 1.8.9) for Estimated Marginal Means analysis.

